# Species diversity increases information flow about predation in bird communities

**DOI:** 10.64898/2026.05.05.722896

**Authors:** Agnishikhe Kumar, Jiahao Wu, Ping Ding, Jakob Bro-Jørgensen, Mylène Dutour, Ari E. Martínez, Xingfeng Si, Qiang Zhang, Eben Goodale

## Abstract

The Biodiversity-Ecosystem Functioning (BEF) literature has shown species diversity to be essential for ecosystem functioning and services. Yet although acquiring information through interspecific networks can impact ecosystem functioning, it is unclear how it is modulated by species diversity. Eliciting vocal responses using predator models across a latitudinal gradient, we first show that the species diversity of birds increases public information about predation both in the low-cost system of mobbing and in the higher-cost system of alarm calls. A similar result was also found across a fragment area gradient for mobbing; this system was then used to test how species diversity affects interspecific information flow in mobbing communities. We set up two BEF playback experiments, manipulating the species richness level of the playback sound files by varying the number of species producing mobbing calls (one, two, four, eight species). In an experiment in which the call rate across treatments was held constant, and only heterospecific responses were counted, increasing species richness of the sound files increased the number of species and individuals responding, the number of calls produced and their frequency range, and decreased latency to call. An experiment in which call rate increased with the addition of species in each treatment showed a similar, but stronger pattern. There was little evidence that the signals of one particular species changed responses. This supports the hypothesis that the species diversity of a community is a key component influencing the quantity and quality of information flow inside it.

## Introduction

The Biodiversity-Ecosystem Functioning (BEF) literature has succeeded over the past three decades to show that species diversity influences ecosystem functioning, often by using manipulative experiments [1, 2]. This literature has shown that differences among species, i.e., complementarity, is a fundamental mechanism that leads to more diverse communities having greater functioning [3, 4]. The results are not only theoretically important, but give further justification stressing the need for biodiversity conservation in the face of global change [3, 5]. While the traditional BEF approach, applied to classical systems like grassland plants, concentrates on the productivity and stability of communities, here we focus on the closely aligned but distinct response variable of information flow, with the goal of applying BEF concepts to behavioral ecology.

Information is a central attribute of living organisms, and can be directly tied to organisms’ fitness in reducing their uncertainty about environmental conditions [6, 7]. Here we will be specifically interested in social information, which is information acquired through interactions with, or observations of, organisms, and is accessible (“public”) to other organisms [8]. Although social information can be acquired from conspecifics, interspecific information (i.e., that sourced from heterospecific individuals) can be especially valuable as it may be acquired with less competition, or species could be complementary in the mechanisms by which they detect, process and transmit information [9–11]. Studies of information flow over the last two decades have shown interspecific information transfer across species – both within and across trophic levels – to be central in making informed decisions [7, 12], influencing individual fitness [13, 14], community assembly [15, 16] and ecosystem functions [17, 18]. Yet despite information being closely linked to the functioning of the ecosystem at varying scales, we do not yet fully understand how the availability and use of public information across species is influenced by species diversity [19].

Here, inspired by the BEF literature, we show the results of a manipulative experiment, demonstrating that the species diversity of information sources increases interspecific information flow within communities (Figure 1A). However, we started the project by first questioning whether species diversity, publicly available information, and interspecific information flow might be related at all, because unlike in classical grassland experiments, animals that are behaviorally interacting can assess the contribution of other species and modify responses without personally contributing. That is, animals can ***eavesdrop*** on information already made public and not contribute themselves, and if this occurred, there might not be any relationship between species diversity and publicly available information, and its flow [20, 21]. To understand how widespread eavesdropping is, a crucial factor is cost: if making signals or cues is costly or risky, all individuals of all species should eavesdrop when publicly available information is sufficient and signals or cues are costly to produce [22].

**Figure 1.**
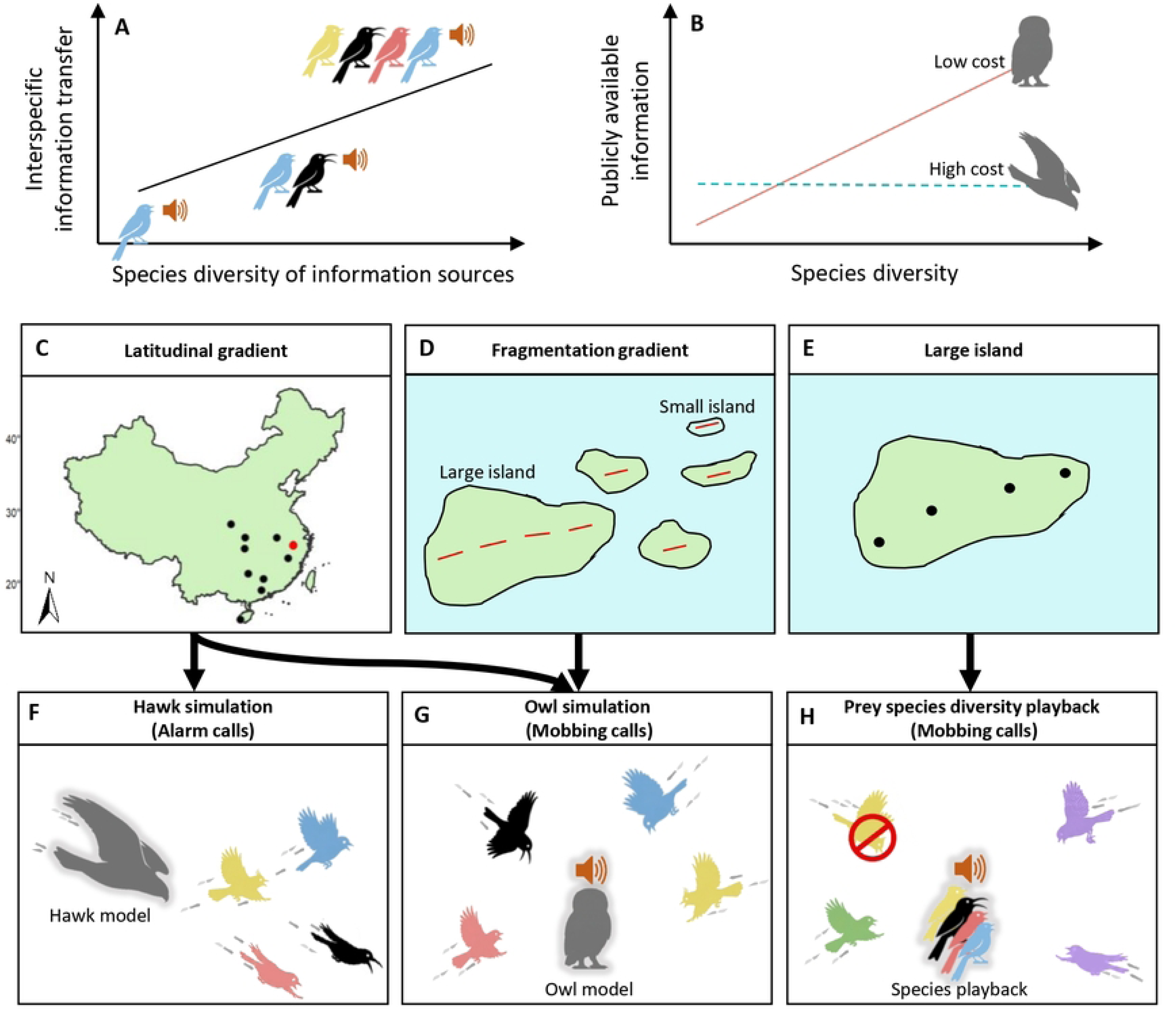
Schematics of species diversity / information relationships and methods across spatial gradients. We hypothesized a positive relationship between species diversity and interspecific information flow (A), and publicly available information about predators, for a low-cost context, but not a high-cost one (B). The effect on availability of public information was assessed over a latitudinal gradient (C, 30 transects in 10 reserves) and over a fragmentation gradient at one reserve, Thousand Island Lake (D, 59 transects on 41 islands). In this work, we elicited alarm calls from mixed-species flocks using hawk models (F) and mobbing calls from bird communities using owl models (G) on the latitudinal gradient, while on the fragmentation one we only elicited mobbing calls. Additionally, on the largest island of the fragmentation gradient (7 sites; E) we conducted manipulative experiments where we played mobbing calls of birds at increasing species richness levels (one, two, four and eight species) to elicit mobbing responses from the bird community (H), excluding the responses by conspecifics.

We take advantage of the complexity of bird communicative signals to focus on cost, because birds have two very contrasting types of signals about predation: alarm calls and mobbing calls [22, 23]. Alarm calls are made in response to high-risk attacks by moving predators and have acoustic qualities to limit their localization by the predator [24]. When alarm calls are made in mixed-species flocks, a common form of avian social organization in forests [25], these calls often lead to flock participants scattering away from the center of the flock into thick vegetation [26]. In contrast, mobbing calls are made to stationary predators that may pose less risk, like an owl in daylight, and the acoustics of their calls optimizes localization, so that more individual prey birds in the community can be recruited towards the caller to harass and ultimately drive the predator away [27]. It therefore can be hypothesized that there should be a strong relationship between species diversity and public information in mobbing, and not in alarm calling (Figure 1B).

We explored the relationship between species diversity and public information availability in high and low-cost contexts on two environmental gradients, a latitudinal gradient (Figures 1C and S1) and a fragmentation gradient (Figures 1D and S2). On the 1650-km latitudinal gradient from tropical to temperate biomes in eastern China, we aimed to compare species diversity and public information relationships in alarm vocalizations in mixed-species flocks, elicited by a sling-shot propelled hawk model (high cost; Figures 1F and S3), to those in mobbing vocalizations elicited by the presentation of an owl model and playback of its calls (low cost; Figures 1G and S3). The fragmentation gradient was located at Thousand Island Lakes (TIL), Zhejiang Province, a subtropical reservoir system that concentrates on changes due to island area [28]; here we did analogous elicitations of mobbing vocalizations only. In both contexts, we measured the following five characteristics of responses: number of species that responded vocally, number of individuals participating (in mobbing only; this could not be estimated for alarms in mixed-species flocks because of their rapidity), call rate [number of calls per minute for all individuals of all species; 29], latency of calls [the length in seconds between when the stimulus was made and the first call; 30], and frequency range (difference between the highest and lowest peak frequencies during the whole trial, including all species’ vocalizations). Call rate is a measure of the quantity of public information, and latency and frequency range indicate its quality. Speed of response is of course important to any response to predation; as to frequency range, as animals are known to respond most to heterospecific alarm calls that share acoustic properties with their own calls [31–33], information from a group performance with a wider frequency range will be transferable to a wider audience [19].

Having demonstrated a species diversity / public information relationship even in the costly context of alarm calls (see Results), we turned to the BEF experiment to investigate how the number of prey species acting as information sources could influence information flow. The large database of mobbing audio recordings from the owl simulation experiment at TIL allowed us to conduct two manipulative playback experiments on the largest island there (Figure 1E). The playback sound files included prey species producing mobbing calls at increasing species richness levels (one, two, four and eight species; Figure 1H) and we measured mobbing responses of birds as before. In the first experiment, the number of calls in the played back sound files increased as a number of species increased (as in reality, the “diversity experiment”), and in the second experiment, the number of calls was controlled to be the same no matter how many species were represented (the “complementarity experiment”).

These two experiments allowed us to distinguish between alternative mechanisms that might produce a positive species diversity and interspecific information flow relationship. First, if there are no differences among species in their signals and cues, more species should stimulate more response in the diversity experiment (Figure 2A), but all treatments should receive the same response in the complementarity experiment (Figure 2B). Here more species simply translate into more individuals and more vocalizations; beyond increasing the quantity of information, this mechanism could affect the quality of information, too, as greater synchrony of calling can lead to higher reliability [34]. Second, if species differ in the characteristics of their signalling, so that the signals they produce are complementary in their characteristics, then the more species that are involved, the more cumulative information will be present [19, 35], which should be seen in both the diversity and complementarity experiments (Figure 2C and D). For instance, as species respond strongly to calls with acoustic characteristics similar to their own, as mentioned above, greater diversity means more receivers can effectively use the information. Third, it could be that the most important factor is not how many individuals or species, but which species are producing information, especially if some species act as crucial sources of information for the whole community [“community informants"; 36, 37]. In this case, known as a selection effect [1, 19], there should be the most response when the community informant species is played back (Figure 2E and F). The selection effect can underlie a positive species diversity and interspecific information flow relationship because a larger species pool has greater probability of including the community informant [1, 19]. In these experiments we also included the calls of a cricket sound as a control, matched to the number of calls in the complementarity experiment, to make sure that response was not merely driven by an intense sound.

**Figure 2.**
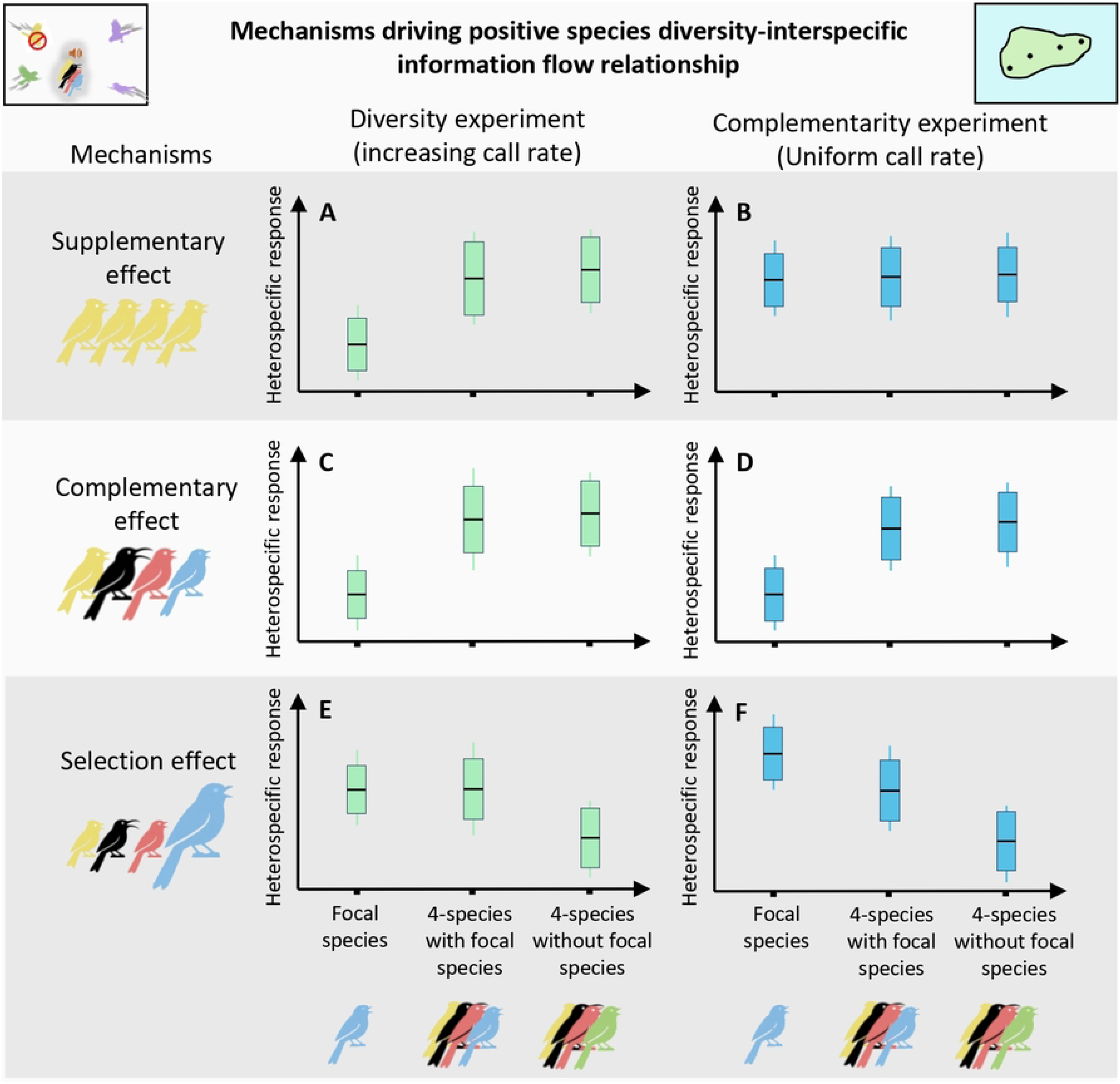
Testing potential mechanisms underlying a positive species diversity and information flow relationship. using the “diversity experiment,” in which call rate increased with the addition of species in each treatment, and the “complementarity experiment”, in which call rate was always held constant. Comparing the heterospecific response to the one-species treatment of each of the focal species with its corresponding four-species treatments, with or without focal species, we could hypothesize: First, for a supplementary effect in which there were no differences among species, the diversity experiment would elicit stronger responses for the four-species treatment with higher call rate (A), while the complementarity experiment would show no difference among treatments (B). Second, if there were complementary differences among species in their signalling, there would be increasing responses for four-species treatments in both experiments (C and D). Third, if the identity of the species were important as in a selection effect, there would be a strong response only in the presence of that species in the treatments in both experiments (E and F).

## Results

### Relationship between species diversity and publicly available information

In over two years of fieldwork, we successfully elicited alarm calls in 79 out of 150 mixed-species flocks we encountered, while also conducted 480 mobbing elicitations. We found a strong relationship between species diversity (of detected species before the trial) and all five of the response variables for mobbing across the latitudinal gradient (Figure 3A-E, Table S1). For alarm calls, the amount of calling was more restrained, as would be expected for the riskier context: whereas there was an average of 409.09 ± 278.33 calls of 4.48 ± 2.38 species per mobbing trial, there were only 4.40 ± 3.16 alarm calls of 0.77 ± 0.85 species per alarm elicitation, and these were made by no more than four species in any trial. Indeed, 60.2% of all alarm calls were made by one species, Huet’s Fulvetta (*Alcippe hueti*), a highly gregarious species, which is the nuclear species that leads mixed-species flocks in the region [38] and also a frequent initiator of mobbing [39]. Nevertheless, there was a positive relationship for alarm calls between species diversity (of the mixed-species flock) and number of species vocalizing, call rate, and frequency range (Figure 3F, G and I). We conclude that even in a situation where calling could involve the ultimate cost – death – there still is evidence that species diversity increases the amount (number of calls) and quality (frequency range) of information. The results of mobbing elicitations at TIL had similar results: species diversity [i.e., species detected before trial, which also strongly correlated with fragment size, 40] significantly influenced all five response variables (Figure S4, Table S2).

**Figure 3.**
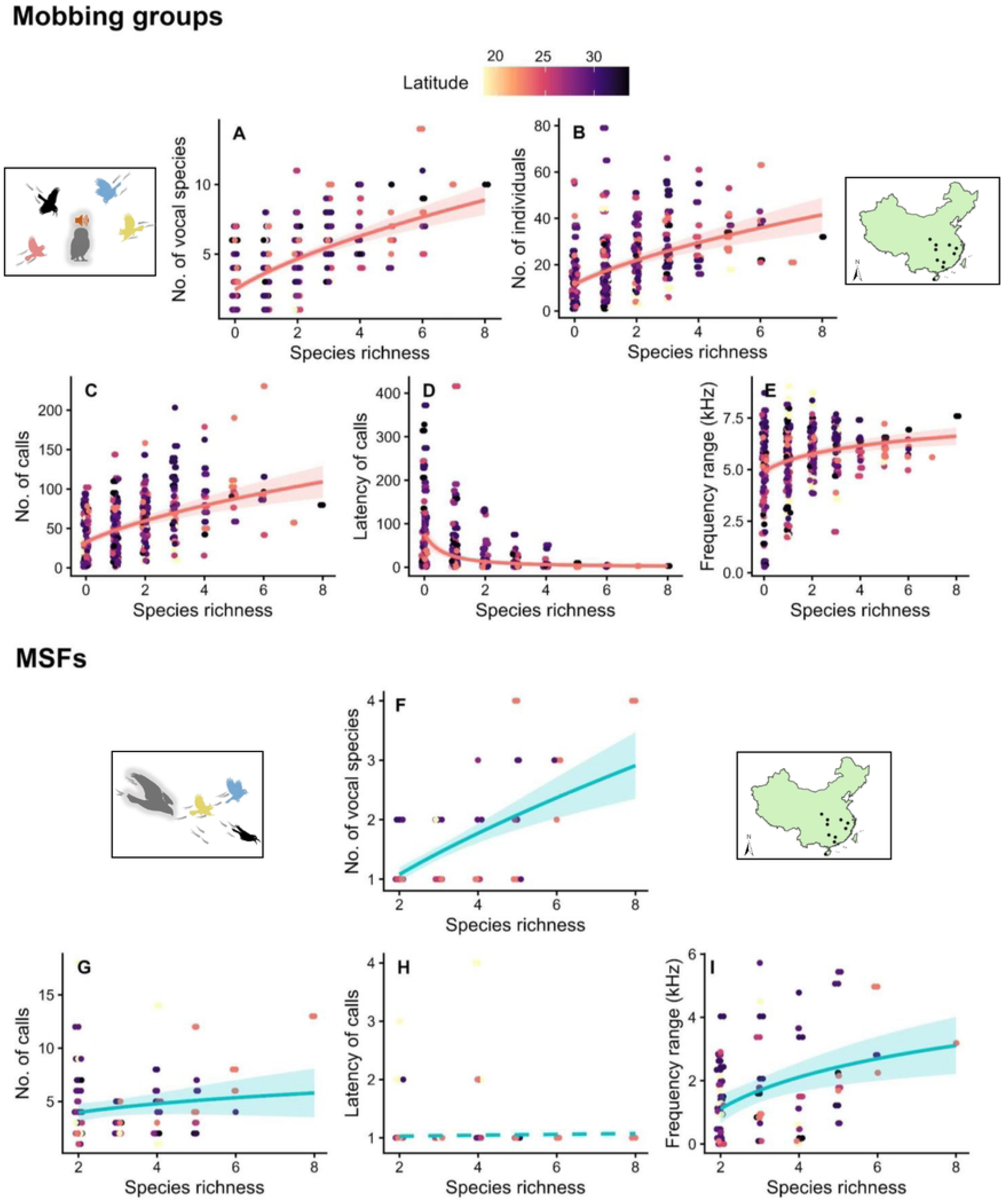
Species richness enhances the quality and quantity of public information along a latitudinal gradient in mobbing assemblages (A-E) and alarm calling within mixed-species flocks (MSFs; F-I). Species richness was assessed as the number of species in the point count before the mobbing elicitation, or the number of species in the flock. The response variables are the number of species that vocalized (A, F), the number of participating individuals (B; this could not be estimated for alarm calls in MSFs), the call rate (per minute; C, G), latency (in s; D, H), and frequency range (kHz; E, I). Solid lines indicate significant predicted values derived from Generalized Linear Mixed Models (GLMMs; Table S1) while dotted lines indicate insignificant effect, and shaded areas represent 95% confidence intervals. Datapoints are colored by latitude (yellow = low latitude; purple = high latitude). Points are jittered along the x-axis to visualize overlapping data.

### Relationship between species diversity and interspecific information flow

We conducted 203 trials of each of the two manipulative experiments; additionally, we conducted seven trials which included the playbacked sounds of the cricket control, which elicited no response. Results of both manipulative experiments showed significant influence of species diversity on all responses, although the responses to the diversity experiment were stronger – i.e., in the amount of variation explained – than to the complementarity experiment (Tables S3 and S4). We focus here on the complementarity experiment (Figure 4; see Figure S5 for results from the diversity experiment). On top (Figure 4A-E), we include all treatments (one through eight species), and responses of all participants (conspecifics – birds responding to the playback of their own species – and heterospecifics), and one can see an increasing but saturating curve, as estimated by GAMMs, for the number of responding species and individuals, the number of calls and the frequency range, and similarly, a decreasing curve for latency. We excluded the eight-species treatment in subsequent analyses with heterospecific responses alone, as the number of available heterospecifics in the community is low because the eight species in the experiment are the most abundant responders. GLMMs only including heterospecific responses also showed significant effects of species richness in enhancing all response variables (Figure 4F-J; Table S4).

**Figure 4.**
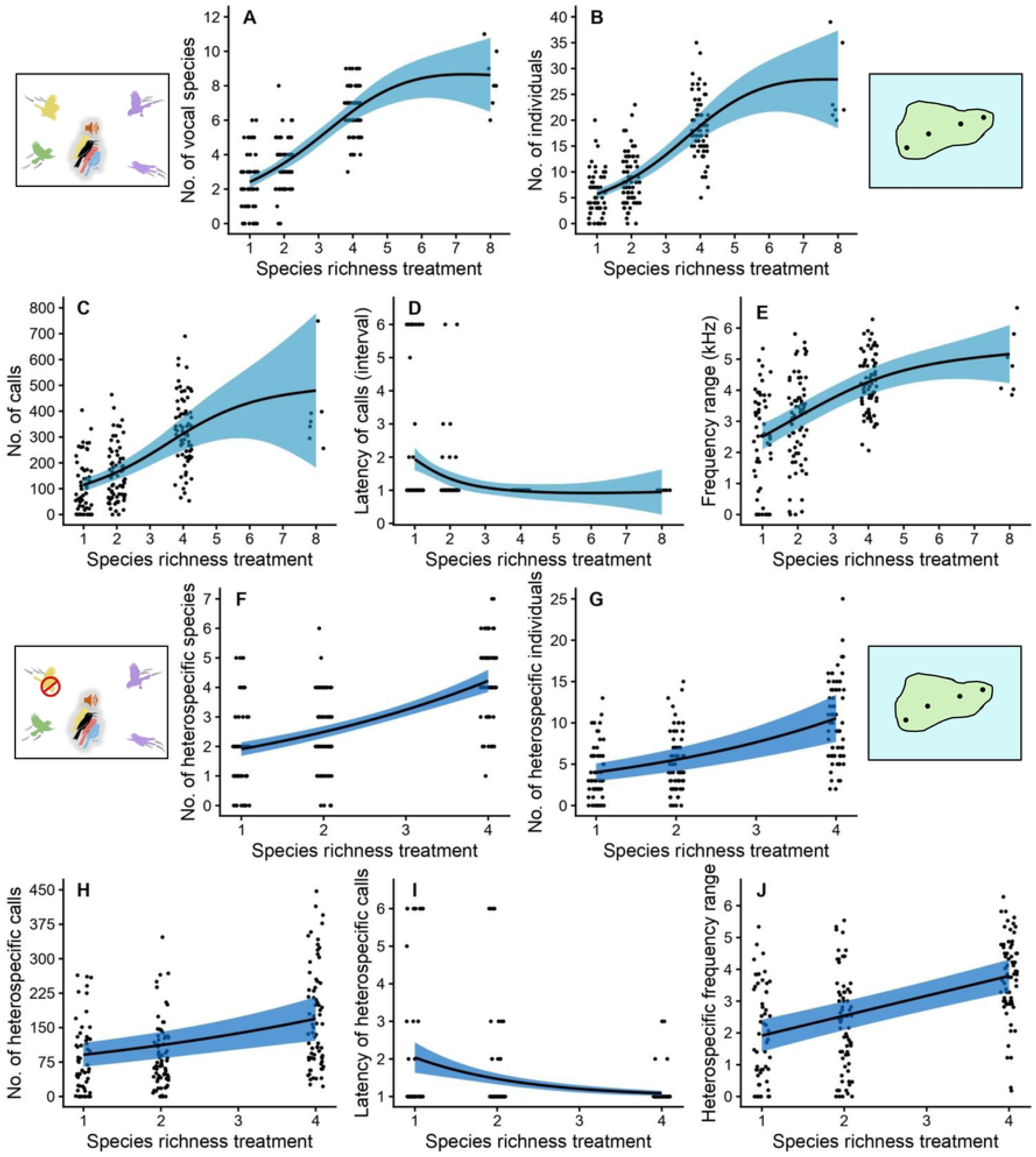
Species richness strengthens interspecific information flow in complementarity experiment. Panels A to E are GAMM models, including the playback treatment with eight species and responses from all species, whereas F to J are the GLMM models without the eight-species treatment and with responses from heterospecific species only. The number of vocal species (A) and participating individuals (B), the number of calls (C), and the frequency range (E) of all species significantly increases with species richness, and the latency of calling declines steadily (D; Table S3). Similarly, the number of heterospecific vocal species (F), participating individuals (G), the number of heterospecific calls (H), and the frequency range (J) also significantly increase with species richness, while the latency of heterospecific calls declining rapidly (I; Table S4). Solid lines indicate predicted values and shaded areas represent 95% confidence intervals. Points are jittered along the x-axis to visualize overlapping data.

To test the mechanisms underlying the species diversity / information flow relationship and the question whether species identity or species richness mattered, we conducted GLMM analyses with Tukey post-hoc tests, comparing heterospecific responses to the one-species treatment of each focal species with those to four-species treatments that a) included that focal species and b) did not. In all 40 models analyzing the diversity experiment (five response variables x eight species; Figures S6-S10; Table S5), one of the four-species treatments had higher responses than the one-species treatment. While the complementary experiment showed weaker results, the general results remained similar: in 32/40 model, one of the four-species treatments had higher responses than the one-species treatment, with the exceptions being 3/8 models from the number of calls, 4/8 from latency and 1/8 from frequency range (Figures 5 and S11-S14; Table S6). A selection effect would show that the inclusion of a key species in playback would elevate response, particularly in the one-species treatment where that species’ vocalizations were not mixed with other species. But none of 80 models (from both experiments) showed highest response to one-species treatment (Figures 5 and S6-S14), and only three models (two for diversity experiment, one for complementarity experiment) showed the four-species treatment with a focal species getting greater response than the four-species treatment without that species (Figures S7C, S8A, and S11C).

**Figure 5.**
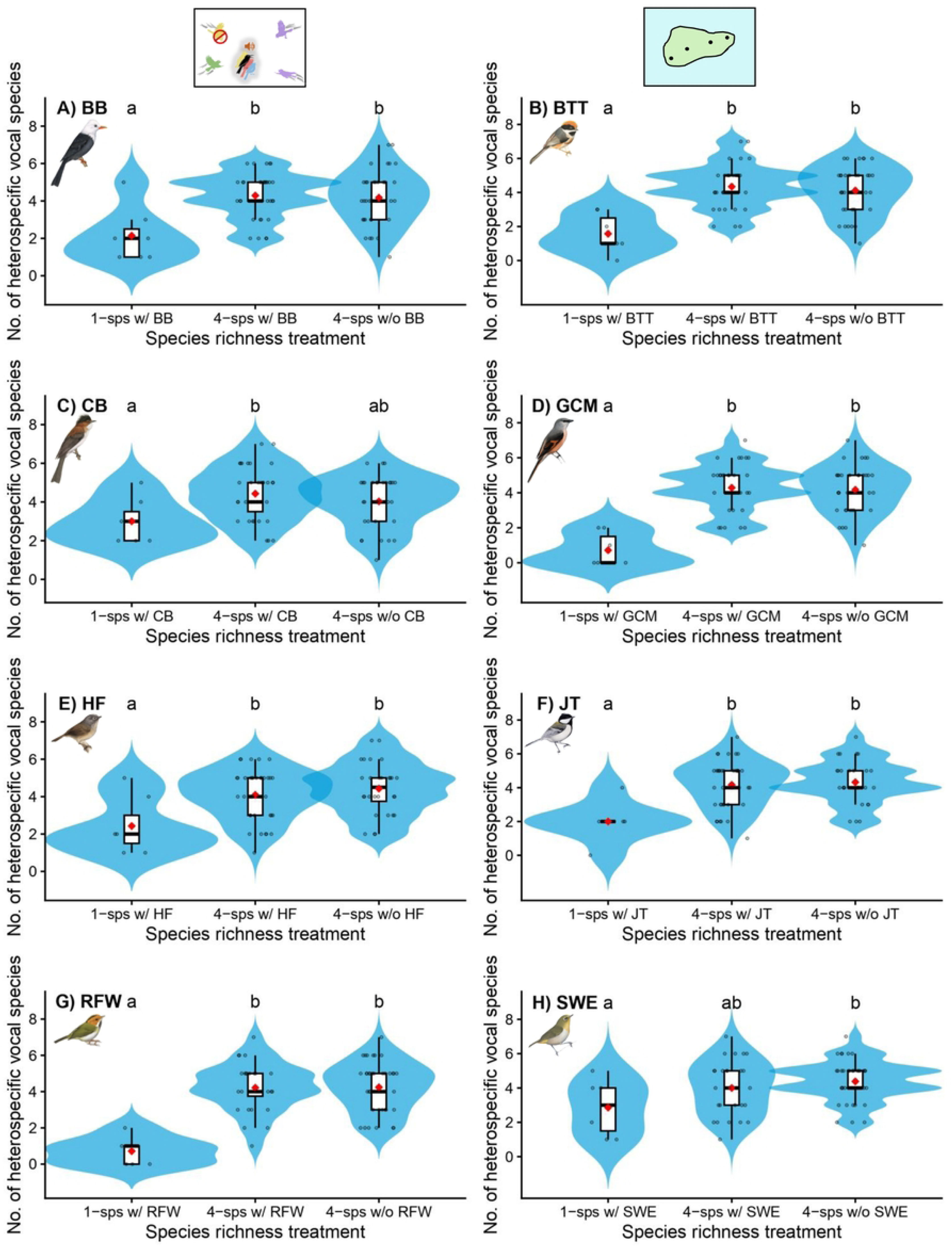
Comparison of the number of vocal species responding to one vs. four-species treatments, for the complementarity experiment. We used GLMM (Table S6), for each of the eight species, to compare how the heterospecific response to the one-species treatment of a focal species varied from four-species treatments, where the focal species was present or absent. Each graph represents one model per species; refer to Table S8 for names of the eight species abbreviated. Black dots are the observed datapoints, the red dot indicates the mean, and the shaded region represents the violin plot. The letters are the results of pairwise comparison between groups from Tukey HSD tests.

## Discussion

### Relationship between species diversity and publicly available information

We worked on the two gradients, latitude and fragmentation, to be able to sample communities of different richness levels. While fragment size strongly influenced species richness of mobbing assemblages (Figure S4), latitude was not a very strong effect on the species richness of either mobs or mixed-species flocks (Figure 3), probably because of the timing of our sampling (to sample more similar communities, we visited reserves from north to south in the non-breeding season, and from south to north in the breeding season). Regardless, public information in mobbing was strongly influenced by species diversity as the majority of species contributed vocalizations [39]. Surprisingly, however, there was also a positive species diversity / public information relationship for alarm calls, even though compared to the mobbing context, the number of calls in a trial was less by two orders of magnitude.

Why would birds continue to alarm call even when other species had already made calls? We can envision two possible reasons. First, perhaps initial alarm calls are unreliable. For example, in a Sri Lankan system, the first species to call is usually the gregarious Orange-billed Babbler *Argya rufescens* [41]. But this species might not be a reliable source of information as it frequently vocalizes to any large or fast-moving object/bird; hence, in a true raptor attack, the calls of other species that come afterwards would be important to emphasize the high threat level. Second, it could be that the cost of calling, although difficult to quantify [42], is actually not as high as is assumed, because to some extent the acoustic characteristics of alarms (i.e. high frequency) might make them difficult to be detected by predators [24]. Yet birds do refrain from calling in some social situations, such as when they are alone or with certain other flock members [43], which suggests some cost, as also reflected in the relatively low number of overall calls that were made. Therefore, we hypothesize that eavesdropping occurs widely in these systems, but a positive species diversity / public information relationship is maintained by the unreliability of initial alarm calls.

### Relationship between species diversity and interspecific information flow

Whereas our data eliciting mobbing and alarm vocalizations with predator models can only address the amount of publicly available information, the manipulative experiments demonstrate that higher species pools translate into greater information usage, and therefore interspecific information transfer. There is little evidence of a selection effect: a selection effect would be suggested by the one-species treatment getting greater response than the four-species treatment, which never happened, or the four-species treatment with the species having greater response than that without it, which happened only in three of 80 models. Instead, both supplementary and complementary mechanisms contribute. The diversity experiment had a stronger response because it includes the supplementary mechanism: as the number of species increases, the number of individuals increases, and each individual produces vocalizations, adding to the collective call rate. In contrast, the complementarity experiment demonstrates that responses increase due to differences among species in their mobbing calls, as the call rate (i.e., simulating individual number) was held constant. Specifically, in the GAMM/GLMM models, latency was less, leading to a quicker response, and the frequency range was broader in response to more played back species, which should translate to a wider audience [19, 32].

Our results are in stronger than earlier findings in this field, which is partially explained by our large-scale design. For example, Igic et al. [44] compared responses to the mobbing vocalizations of two individuals of two species to that of two individuals of one species, and found no difference. But responses were only observed for a species of magpie, rather than the community level as tested here, and the differences in species diversity (one vs. two) was narrower than what tested here (one to eight). Todd Freeberg and colleagues have investigated the effect of species diversity of mixed-species flocks on the decisions and communication of flock participants. While species diversity did increase problem solving at a novel feeder [45], the effects on communication were weak [46, 47]. Yet this temperate flock system also has a limited amount of diversity (tests mostly compared one versus two species). Our results may show the benefits of working on subtropical and tropical systems when exploring the effect of species diversity.

As with other examples of BEF studies that illustrate the importance of species diversity on ecosystem functioning and services such as pollination [48], seed dispersal [49] and pest control [50], our result of a positive information-species diversity relationship helps justify biodiversity conservation. Our result adds to the evidence that in human-modified landscapes where species diversity is threatened, the remaining animals face problems associated with having few individuals / species present [40, 51]. If social information is unavailable or limited, animals might make poorly informed decisions, for instance, either remaining uninformed of predators, or showing weaker response, whether that be individual (e.g., escape) or collective (e.g., confusing or driving away a predator through mobbing) behavior. Further, individuals may turn to personally collected information, which may prove to be energetically costly and time-consuming, or even expose the animal to predation risk [52, 53].

Although we have built this argument about heterospecific information about predation in birds as our model system, we emphasise that the importance of species diversity for interspecific information transfer can be applied to other behaviors such as selection of resources or habitats, and could involve a wide range of animals including invertebrates, interacting both within and across trophic levels [17, 19]. For example, migratory species could eavesdrop on cues from communities to select high quality sites, or animals can hone in on the signals and cues of heterospecifics at shared resources [54–57]. Situations in which the information can be manipulated, such as using playback here, or chemical cues/signals in invertebrate systems [58], or herbivore-induced plant volatiles in multi-trophic level interaction networks [56, 59], may be particularly useful to understand the theoretical and conservation implications of interspecific information transfer.

## Materials and Methods

### Ethical statement

All aspects of this study received ethical permission from Xi’an Jiaotong-Liverpool University (ER-UEC-11000092420230130213130). The fieldwork was also approved of by the relevant authorities for the ten different nature reserves.

### Predator simulations experiments testing the species diversity / publicly available information relationship

#### Study sites for the latitudinal gradient

For the latitudinal fieldwork, we selected ten nature reserves in China (Figure S1). All the reserves were within the range of the Collared Owlet, *Taenioptynx brodiei*, an owlet that can attack birds; several *Accipiter* bird-eating hawk species are also widespread in this region. At each reserve, we set up three 2-km transects spaced at least 500 m apart from each other. Other than the two extremes of the network (Hainan, tropical; Shaanxi, temperate), most of these nature reserves are subtropical and represent relatively mature secondary forest that is currently well-protected (Table S7). We visited this network in the breeding (May-August) and non-breeding season (November-February) twice per year across two annual cycles (2023-2024 and 2024-2025). While mobbing data was collected in both seasons, alarm calling in mixed-species flocks data was taken only during the non-breeding season, as subtropical flock systems can be unstable in the breeding period [60]. To make the data most similar between reserves, we sampled them from south to north in the breeding season, and from north to south in the non-breeding season.

#### Study site for the fragmentation gradient

The fragmentation gradient was studied at Thousand Island Lake (TIL), Zhejiang Province, which was one of the 10 reserves in the latitudinal study, and serves as one of the foremost platforms for the study of habitat fragmentation and island biogeography in China [28]. Here we used an existing network of 59 transects laid across 41 islands (Figure S2), with the number of transects per island proportional to the log area of the island. These transects are generally 400 m long, except for a few that are shorter to fit in very small islands (so on average they are 321.19 ± 82.63 m in length). The fragmentation gradient was sampled for mobbing data in the non-breeding (December 2023 to January 2024) and breeding season (May 2024 to June 2024).

#### Hawk predator simulation experiment eliciting alarm calling behavior

To elicit alarm calls in mixed-species flocks, we first walked the transects on the latitudinal gradient in search of flocks. On encountering a mixed-species flock, we ensured the flock had habituated to our presence, and collected the data on the individuals and species in the flock (Figure S3A). We then audio recorded 1 min of baseline activity (Figure S3B), with a MixPre-3 II recorder and an Audio-Technica AT4022 omnidirectional microphone embedded in a Telinga parabola (producing .wav files with 48 kHz sampling rate and 24-bit depth). We then fired a 20-cm slingshot-propelled plastic hawk model to a height of ∼10 m and a distance of ∼20 m to one side of the front of the flock (Figure S3C). Recording continued until all alarm calls stopped or for 1 min after the model was shot. For the mixed-species flock context, because alarms were so rapid and short, it was not possible to estimate the number of individuals involved.

#### Owl predator simulation experiment eliciting mobbing behavior

The elicitation of mobbing vocalizations in both the latitudinal and fragmentation gradient was done similarly, and analogously to the elicitation of alarm calls. At points on the transect (at both ends of transects for the fragmentation gradient, except for one transect so short that it had only one playback site; at 250 m intervals for the latitudinal gradient), we first did a fixed radius point count (7-min for fragmentation gradient and 5-min for latitudinal gradient; Figure S3D), noting all species seen or heard within 30 m, for each trial. Then, the recordings started with 1 min of baseline activity (Figure S3E), followed by 7 min of the playback experiment (Figure S3F). During the playback experiment, we played a randomly chosen playback tape of Collared Owlet (of the four playback tapes prepared from high signal-to-noise ratio recordings on Xeno-Canto.org) on a JBL Flip 6 speaker, broadcast at 90 dB at 1 m. These calls played for 30 s, followed by 30 s of silence and were repeated seven times. Individuals were considered to have participated in mobbing if they approached the speaker within 30 m during the playback period, and exhibited mobbing behavior [rapid head-scanning or perch-changing, mobbing vocalizations; 61].

#### Measuring responses to the predator simulation experiments

We measured similar responses for both the predator simulation experiments on the different gradients. A species participating in a trial was said to have vocalized if it produced alarm call or mobbing call, respectively. An alarm call or mobbing call was a vocalization that started after the stimulus or represented a change in vocalization type, checking that the same vocal type was also used in the majority of other trials for the species [41, 60]. In practice, the vocal types used aligned with those found on Xeno-Canto in predator contexts. We measured the five aspects of responses, with the first two indicating alarm/mobbing participation and the latter three indicating metrics of information: 1) the number of species that responded vocally, 2) number of individuals that participated in the trial (estimated only for the mobbing context), 3) call rate (calls per min), 4) latency of calls (time in s) and 5) frequency range (kHz). All responses except the number of participating individuals (noted during the experiment) were measured in Raven Pro acoustic software (version 1.6) using the spectrograms of the recording with Hann window and Fourier transformation (FFT) size of 512 samples settings.

#### Analysis of the vocalizations elicited to the predator simulation experiments

We used general linear mixed models (GLMM) in the “lme4” package to analyse the predator simulation data. We modelled five models, one for each of the five response variables, and used species richness as a fixed predictor and transect identity as a random factor. The model distributions were chosen based on the residual diagnostics, testing for normality, dispersion, heteroscedasticity, zero-inflation and outliers using “DHARMa” package. We chose Gaussian distribution for continuous data (applying log transformation or log link when appropriate), Poisson error distribution (log link) for count data, Conway-Maxwell-Poisson distribution (aka Compois, log linked) for under-dispersed data, negative binomial (NB, log linked) distribution for over-dispersed data, Zero-inflated NB distribution (log linked) for data with zero inflation, and Truncated NB distribution (log linked) and Truncated Compois distribution (log linked) for right skewed data with non-zeros.

### Manipulative playback experiments testing the species diversity / interspecific information flow relationship and mechanisms

#### Study site

The two manipulative experiments – the diversity and complementarity experiments – were conducted at seven sites at least 400 m apart from each other on the largest island fragment (1289.23 ha) in TIL.

#### Experimental design

The eight most frequently mobbing species from the owl predator simulation experiment on larger islands in TIL (Table S8) were used as our species pool to construct the playback sounds for the diversity and complementarity experiments. We designed four treatments with species producing mobbing calls at increasing species richness levels, i.e., one, two, four, and eight species (Table S9). The complementarity experiment made sure that all treatments had 104 calls per minute (the average for all species across all predator simulation trials on the larger islands of TIL), whereas for the diversity experiment, we used the average number of calls that were made by a species (Table S8), and hence the calls per min increased with the addition of species. For both experiments, the one-species treatment had eight sub-treatments (one per species in the species pool; Table S9). The two-species and four-species treatments had 10 sub-treatments each, randomly chosen from all possible combinations in the species pool (see Table S9). The eight-species treatment was represented with a single sub-treatment including all eight species. We used a single sub-treatment of Japanese Burrowing Cricket sounds (*Velarifictorus micado*, present at the study site, personal observation, AK) as a control (matched to the call rate of the complementarity experiment). In total, 29 sub-treatments from four treatments of both experiments, and one control were played at each of the seven sites (29 sub-treatments per experiment x 7 sites = 203 trials; 203 trials x 2 experiments + 1 control x 7 sites = 413 trials total).

#### Playback sound files

Each sub-treatment was represented by three exemplar playback sound files, with each sound file made from a different set of recordings. Each set contained eight clean recordings per species recorded from different clusters of islands to ensure independence among sound files. While preparing the recordings, we removed low frequency noise, background noise, calls of non-focal birds and standardized volume of focal species to be similar to field observations (Table S8). The sound files were constructed on the sound editing software Veed (veed.io) with the calls from the recordings arranged together in a randomized order to mimic a natural mobbing assemblage. Sound files were 5 min long with alternating 40 s of playback period and 20 s silent periods (to help evaluate responses), repeated five times. As the manipulative experiments used two speakers to simulate the presence of multiple individuals (see trial protocol), we constructed two replicates for each sound file so that the speakers did not play in unison. Accordingly, we first constructed a sound file containing half of the total calls (104/2 calls for the complementarity experiment, and n/2 calls for the diversity experiment, where n is the total number of calls per sub-treatment). We then reversed the order of the two 20-s halves of the 40-s call sequence in that sound file, resulting in two replicates that were played at once.

#### Trial protocol

The trial and its recording were similar to the owl predator simulation experiment. For every trial at each site, we first conducted a 5-min point count (Figure S3G), followed by 1-min recording of the baseline activity (Figure S3H), and 5 min of the playback period (Figure S3I). For the playback, we played a randomly selected playback sound file at two JBL speakers spaced 8-10 m away from each other and recorded the mobbing response.

#### Measuring responses and analysis

Analysis for the complementarity and diversity experiments were similar to that for the predator simulation data. However, in an effort to blind the dataset, here we removed the parts of the recordings in which the playback could be heard and renamed the sound files. The data were only afterwards filtered for heterospecifics. This blinding procedure could not be used for the number of participating individuals, which used field data, and the frequency range, which was difficult to score without knowing beforehand which species were heterospecific. As we extracted data from the silence periods, we express the intensity of calling as the total number of calls produced during silent periods and latency as the interval at which the participants first called (first to sixth; with the sixth indicating no response) rather than the time in s. To assess the effect of species richness on mobbing responses from all species/individuals (including heterospecifics – i.e., those not on the sound file played back – and conspecifics), we used generalized additive mixed models (GAMM; “mgcv” package) that were estimated with restricted maximum likelihood. We used species richness as the main predictor with a smoothing term (k = 4) and site as a random predictor. To further assess the effect of species richness on heterospecific responses alone (one-species to four-species data), we used GLMMs. To test the mechanisms and differentiate whether the heterospecific responses were driven by species richness or species identity, we compared how the one-species treatment of the focal species varied from four-species treatments, with and without the focal species using GLMMs and Tukey HSD tests.

## Acknowledgements

We thank Xin’an River Ecological Development Group Corporation, Chun’an Forestry Bureau and the Thousand Island Lake National Forest Park, and the administration of the other nine reserves, for their support of the fieldwork. We are grateful to Zijuan Cao and Chinese National Geography Intellectual Property Co. Ltd for the bird illustrations, and their artists Xiaodong Li, Jing Qian, Shaochong Peng, Liang Su, Bai Xiao, and Qiuyang Zheng. We appreciate the help of Yuehan Dou, Juan Li, Li Li, Yaoqi Li, Mike Speed, Lingyun Xiao, and Yi Zou for their helpful comments as the project developed. We acknowledge the use of large language models, such as ChatGPT, for their assistance in the development of initial coding for some parts of the analysis and for the illustrations of bird silhouettes in the figures.

## Author contributions

**Conceptualization:** E.G., A.E.M., X.S., and Q.Z.

**Methodology:** A.K., J.W., M.D., and E.G.

**Formal Analysis:** A.K., and J.W.

**Investigation:** A.K., and J.W.

**Visualization:** A.K., J.W., and E.G.

**Funding acquisition:** E.G., P.D., X.S., and Q.Z.

**Project administration:** E.G., P.D., X.S., and Q.Z.

**Supervision:** E.G., and J.B-J.

**Writing – original draft:** A.K., J.W., and E.G.

**Writing – review & editing:** A.K., J.W., P.D., J.B-J., M.D., A.E.M., X.S., Q.Z., and E.G.

## Data availability statement

The data and the code supporting the findings of the study are available on Figshare at https://figshare.com/s/dde74c73382b6b39472c and https://figshare.com/s/479b99310b605c3609f9

## Supplemental Information Figure and Table captions

Document S1. Tables S1-9 and Figures S1-14

Table S1. GLMM results for the latitudinal gradient.

Table S2. GLMM results for the fragmentation gradient.

Table S3. GAMM results from the manipulative experiments.

Table S4. GLMM results from the manipulative experiments.

Table S5. Comparison of responses to one vs four-species treatments, for the diversity experiment.

Table S6. Comparison of responses to one vs four-species treatments, for the complementarity experiment.

Table S7. The national nature reserves sampled across the latitudinal gradient.

Table S8. The characteristics of the mobbing calls of the eight most frequent species.

Table S9. The treatments and sub-treatments in the manipulative experiment.

Figure S1. The map of the latitudinal gradient in China.

Figure S2. The map of the fragmentation gradient.

Figure S3. The protocol of the predation simulation and the manipulative experiments .

Figure S4. The species richness / public information relationship along the fragmentation gradient.

Figure S5. The species richness / information flow relationship in the diversity experiment.

Figure S6. Comparison of the number of vocal species responding to one vs. four-species treatments, for the diversity experiment.

Figure S7. Comparison of the number of individuals responding to one vs. four-species treatments, for the diversity experiment.

Figure S8. Comparison of the number of calls to one vs. four-species treatments, for the diversity experiment.

Figure S9. Comparison of the latency of calling to one vs. four-species treatments, for the diversity experiment.

Figure S10. Comparison of the frequency range in one vs. four-species treatments, for the diversity experiment.

Figure S11. Comparison of the number of individuals responding to one vs. four-species treatments, for the complementarity experiment.

Figure S12. Comparison of the number of calls to one vs. four-species treatments, for the complementarity experiment.

Figure S13. Comparison of the latency of calling to one vs. four-species treatments, for the complementarity experiment.

Figure S14. Comparison of the frequency range in one vs. four-species treatments, for the complementarity experiment.

## References

1. Tilman D, Isbell F, Cowles JM. Biodiversity and ecosystem functioning. Annual Review of Ecology, Evolution, and Systematics. 2014;45:471–93. doi: 10.1146/annurev-ecolsys-120213-091917.

2. Hooper DU, Chapin FS, Ewel JJ, Hector A, Inchausti P, Lavorel S, et al. Effects of biodiversity on ecosystem functioning: A consensus of current knowledge. Ecological Monographs. 2005;75(1):3–35. doi: 10.1890/04-0922.

3. Cardinale BJ, Duffy JE, Gonzalez A, Hooper DU, Perrings C, Venail P, et al. Biodiversity loss and its impact on humanity. Nature. 2012;486(7401):59–67. doi: 10.1038/nature11148.

4. Loreau M, Hector A. Partitioning selection and complementarity in biodiversity experiments. Nature. 2001;412(6842):72–6. doi: 10.1038/35083573.

5. Dee LE, De Lara M, Costello C, Gaines SD. To what extent can ecosystem services motivate protecting biodiversity? Ecology Letters. 2017;20(8):935–46. doi: 10.1111/ele.12790. PubMed PMID: WOS:000405917500001.

6. Dall SR, Giraldeau L-A, Olsson O, McNamara JM, Stephens DW. Information and its use by animals in evolutionary ecology. Trends in Ecology & Evolution. 2005;20(4):187–93. doi: 10.1016/j.tree.2005.01.010.

7. Schmidt KA, Dall SRX, van Gils JA. The ecology of information: An overview on the ecological significance of making informed decisions. Oikos. 2010;119:304–16. doi: 10.1111/j.1600-0706.2009.17573.x.

8. Wagner RH, Danchin É. A taxonomy of biological information. Oikos. 2010;119(2):203-doi: 10.1111/j.1600-0706.2009.17315.x.

9. Seppänen J-T, Forsman JT, Mönkkönen M, Thomson RL. Social information use is a process across time, space and ecology, reaching heterospecifics. Ecology 2007;88:1622–33. doi: 10.1890/06-1757.1.

10. Goodale E, Sridhar H, Sieving K, Bangal P, Colorado Zuluaga G, Farine D, et al. Mixed company: A framework for understanding the composition and organization of mixed-species animal groups. Biological Reviews. 2020;95:889–910. doi: 10.1111/brv.12591.

11. Turner CR, Spike M, Magrath RD. The evolution of eavesdropping on heterospecific alarm calls: Relevance, reliability, and personal information. Ecology and Evolution. 2023;13:e10272. doi: 10.1002/ece3.10272.

12. Farine DR, Aplin LM, Sheldon BC, Hoppitt W. Interspecific social networks promote information transmission in wild songbirds. Proceedings of the Royal Society B. 2015;282:20142804. doi: 10.1098/rspb.2014.2804.

13. Dolby AS, Grubb Jr TC. Benefits to satellite members in mixed-species foraging groups: an experimental analysis. Animal behaviour. 1998;56(2):501–9. doi: 10.1006/anbe.1998.0808.

14. Forsman J, Seppänen J-T, Mönkkönen M. Positive fitness consequences of interspecific interaction with a potential competitor. Proceedings of the Royal Society of London Series B: Biological Sciences. 2002;269(1500):1619–23. doi: 10.1098/rspb.2002.2065.

15. Goodale E, Beauchamp G, Magrath R, Nieh JC, Ruxton GD. Interspecific information transfer influences animal community structure. Trends in Ecology and Evolution. 2010;25:354–61. doi: 10.1016/j.tree.2010.01.002.

16. Gil MA, Hein AM, Spiegel O, Baskett ML, Sih A. Social information links individual behavior to population and community dynamics. Trends in Ecology and Evolution. 2018;33(7):535–48. doi: 10.1016/j.tree.2018.04.010. PubMed PMID: WOS:000438464300012.

17. Brose U, Hirt MR, Ryser R, Rosenbaum B, Berti E, Gauzens B, et al. Embedding information flows within ecological networks. Nature Ecology and Evolution. 2025:1–12. doi: 10.1038/s41559-025-02670-2.

18. Gil MA, Hein AM. Social interactions among grazing reef fish drive material flux in a coral reef ecosystem. Proceedings of the National Academy of Sciences. 2017;114(18):4703–8. doi: 10.1073/pnas.1615652114.

19. Goodale E, Magrath RD. Species diversity and interspecific information flow. Biological Reviews. 2024;99:999–1014. doi: 10.1111/brv.13055.

20. Ridley AR, Wiley EM, Thompson AM. The ecological benefits of interceptive eavesdropping. Functional Ecology. 2014;28(1):197–205. doi: 10.1111/1365-2435.12153.

21. Sullivan K. Selective alarm calling by downy woodpeckers in mixed-species flocks. The Auk. 1985:184–7. doi: 10.2307/4086843.

22. Magrath RD, Haff TM, Fallow PM, Radford AN. Eavesdropping on heterospecific alarm calls: From mechanisms to consequences. Biological Reviews. 2015;90(2):560–86. doi: 10.1111/brv.12122. PubMed PMID: WOS:000352818700011.

23. Marler P. Characteristics of some animal calls. Nature. 1955;176(4470):6–8. doi: 10.1038/176006a0.

24. Klump G, Kretzschmar E, Curio E. The hearing of an avian predator and its avian prey. Behavioral Ecology and Sociobiology. 1986;18(5):317–23. doi: 10.1007/BF00299662.

25. Goodale E, Beauchamp G, Ruxton GD. Mixed-species groups of animals: behavior, community structure, and conservation: Academic Press; 2017.

26. Gaddis P. Mixed flocks, accipiters, and antipredator behavior. Condor. 1980;82(3):20.

27. Carlson NV, Freeberg TM, Goodale E, Theo AH. Mixed-species groups and aggregations: shaping ecological and behavioural patterns and processes. Philosophical Transactions of the Royal Society B: Biological Sciences. 2023;378(1878). doi: 10.1098/rstb.2022.0093.

28. Si X, Jin T, Li W, Ren P, Wu Q, Zeng D, et al. TIL20: A review of island biogeography and habitat fragmentation studies on subtropical reservoir islands of Thousand Island Lake, China. Zoological Research: Diversity and Conservation. 2024;1(2):89–105. doi: 10.24272/j.issn.2097-3772.2024.001.

29. Fernández GJ, Dutour M, Carro ME. Information transfer during mobbing: Call rate is more important than the number of callers in a southern temperate passerine. Behavioral Ecology and Sociobiology. 2023;77(7):78. doi: 10.1007/s00265-023-03357-z.

30. McLachlan JR, Magrath RD. Speedy revelations: How alarm calls can convey rapid, reliable information about urgent danger. Proceedings of the Royal Society B. 2020;287(1921):20192772. doi: 10.1098/rspb.2019.2772.

31. Johnson FR, McNaughton EJ, Shelley CD, Blumstein DT. Mechanisms of heterospecific recognition in avian mobbing calls. Australian Journal of Zoology. 2004;51(6):577–85. doi: 10.1071/ZO03031.

32. Fallow PM, Gardner JL, Magrath RD. Sound familiar? Acoustic similarity provokes responses to unfamiliar heterospecific alarm calls. Behavioral Ecology. 2011;22(2):401–10. doi: 10.1093/beheco/arq221.

33. Meise K, Franks DW, Bro-Jorgensen J. Multiple adaptive and non-adaptive processes determine responsiveness to heterospecific alarm calls in African savannah herbivores. Proceedings of the Royal Society B. 2018;285:20172676. doi: 10.1098/rspb.2017.2676. PubMed PMID: WOS:000438492800001.

34. Wolf M, Kurvers R, Ward AJW, Krause S, Krause J. Accurate decisions in an uncertain world: Collective cognition increases true positives while decreasing false positives. Proceedings of the Royal Society B. 2013;280(1756):20122777. doi: 10.1098/rspb.2012.277710.1098/rspb.2012.2777. PubMed PMID: WOS:000315461500015.

35. Carlson NV, Healy SD, Templeton CN. A comparative study of how British tits encode predator threat in their mobbing calls. Animal Behaviour. 2017;125:77–92. doi: 10.1016/j.anbehav.2017.01.011. PubMed PMID: WOS:000398050900014.

36. Hetrick SA, Sieving KE. Antipredator calls of tufted titmice and interspecific transfer of encoded threat information. Behavioral Ecology. 2012;23(1):83–92. doi: 10.1093/beheco/arr16010.1093/beheco/arr160. PubMed PMID: WOS:000298386500011.

37. Meise K, Franks DW, Bro-Jørgensen J. Alarm communication networks as a driver of community structure in African savannah herbivores. Ecology Letters. 2020;23(2):293–304. doi: 10.1111/ele.13432.

38. Zhou L, Peabotuwage I, Gu H, Jiang D, Jiang A, Zhang M, et al. The response of mixed-species flocks to anthropogenic disturbance and elevational change in southwest China. Condor. 2019;121:duz028. doi: 10.1093/condor/duz028.

39. Wu J, Shen, Y., Bro-Jorgensen, J., Zhang, Q., and Goodale, E. Multiple community informants enhance public information in a transition zone. Integrative Zoology. In review.

40. Kumar A, Ding, P., Bro-Jørgensen, J., Mammides, C., Martínez, A. E., Si, X., Goodale, E. . Habitat fragmentation reduces the quantity and quality of information about predators in avian communities. . Journal of Animal Ecology. In review.

41. Goodale E, Kotagama SW. Alarm calling in Sri Lankan mixed-species bird flocks. Auk. 2005;122:108–20. doi: 10.1093/auk/122.1.108.

42. Sherman PW. Nepotism and the evolution of alarm calls. Science. 1977;197:1246–53.

43. Krama T, Krams R, Elferts D, Sieving KE, Krams IA. Selective selfishness in alarm calling behaviour by some members of wintering mixed-species groups of crested tits and willow tits. Philosophical Transactions of the Royal Society B. 2023;378(1878):20220102. doi: 10.1098/rstb.2022.0102.

44. Igic B, Ratnayake CP, Radford AN, Magrath RD. Eavesdropping magpies respond to the number of heterospecifics giving alarm calls but not the number of species calling. Animal Behaviour. 2019;148:133–43. doi: 10.1016/j.anbehav.2018.12.012. PubMed PMID: WOS:000457630500014.

45. Freeberg TM, Eppert SK, Sieving KE, Lucas JR. Diversity in mixed species groups improves success in a novel feeder test in a wild songbird community. Scientific Reports. 2017;7:43014. doi: 10.1038/srep43014. PubMed PMID: WOS:000394767700001.

46. Brooks HJ, Freeberg TM. Single-species and multi-species playbacks elicit asymmetrical responses within mixed-species chickadee, titmouse, and nuthatch flocks. Ethology. 2024;130(6):e13459. doi: 10.1111/eth.13459.

47. Frazier EK, Selman ZA, Price CA, Papeş M, Freeberg TM. Mixed-Species Flock Diversity and Habitat Density Are Associated with Antipredator Behavior in Songbirds. Diversity. 2025;17(5):363. doi: 10.3390/d17050363.

48. Hadley AS, Betts MG. The effects of landscape fragmentation on pollination dynamics: absence of evidence not evidence of absence. Biological Reviews. 2012;87(3):526–44. doi: 10.1111/j.1469-185X.2011.00205.x.

49. Zhu C, Dalsgaard B, Li W, Kaiser-Bunbury CN, Simmons BI, Ren P, et al. Interconnecting fragmented forests: Small and mobile birds are cornerstones in the plant–frugivore meta-network. Proceedings of the National Academy of Sciences. 2025;122(7):e2415846122. doi: 10.1073/pnas.2415846122.

50. Bregman TP, Sekercioglu CH, Tobias JA. Global patterns and predictors of bird species responses to forest fragmentation: implications for ecosystem function and conservation. Biological Conservation. 2014;169:372–83. doi: 10.1016/j.biocon.2013.11.024.

51. Schmidt KA, Johansson J, Betts MG. Information-mediated Allee effects in breeding habitat selection. American Naturalist. 2015;186(6):E162-E71. doi: 10.1086/683659. PubMed PMID: WOS:000365307700002.

52. Laland KN. Social learning strategies. Learning and Behavior. 2004;32(1):4-14. doi: 10.3758/BF03196002.

53. Webster MM, Laland KN. Social learning strategies and predation risk: Minnows copy only when using private information would be costly. Proceedings of the Royal Society B: Biological Sciences. 2008;275(1653):2869–76. doi: 10.1098/rspb.2008.0817.

54. Sebastián-González E, Morales-Reyes Z, Botella F, Naves-Alegre L, Pérez-García JM, Mateo-Tomás P, et al. Network structure of vertebrate scavenger assemblages at the global scale: drivers and ecosystem functioning implications. Ecography. 2020;43(8):1143–55. doi: 10.1111/ecog.05083.

55. Gu H, Chen J, Ewing H, Liu X, Zhao J, Goodale E. Heterospecific attraction to the vocalizations of birds in mass-fruiting trees. Behavioral Ecology and Sociobiology. 2017;71(5):82. doi: 10.1007/s00265-017-2312-6. PubMed PMID: WOS:000401148300006.

56. Kigathi RN, Weisser WW, Reichelt M, Gershenzon J, Unsicker SB. Plant volatile emission depends on the species composition of the neighboring plant community. BMC Plant Biology. 2019;19(1):58. doi: 10.1186/s12870-018-1541-9.

57. Slaa EJ, Hughes WOH. Local enhancement, local inhibition, eavesdropping, and the parasitism of social insect communication. In: Hrncir M, Jarau S, editors. Food Exploitation by Social Insects. Boca Raton, FL, USA: CRC Press; 2009. p. 147-64.

58. Nieh JC, Barreto LS, Contrera FA, Imperatriz–Fonseca VL. Olfactory eavesdropping by a competitively foraging stingless bee, Trigona spinipes. Proceedings of the Royal Society of London Series B: Biological Sciences. 2004;271(1548):1633–40. doi: 10.1098/rspb.2004.2717.

59. Amo L, Jansen JJ, van Dam NM, Dicke M, Visser ME. Birds exploit herbivore-induced plant volatiles to locate herbivorous prey. Ecology Letters. 2013;16(11):1348–55. doi: 10.1111/ele.12177.

60. Jiang D, Sieving KE, Meaux E, Goodale E. Seasonal changes in mixed-species bird flocks and antipredator information. Ecology and Evolution. 2020;10:5368–82. doi: 10.1002/ece3.6280.

61. Carlson NV, Griesser M. Mobbing in animals: A thorough review and proposed future directions. Advances in the Study of Behavior. 2022;(54):1–41. doi: 10.1016/bs.asb.2022.01.003.

